# Differences in components of reproductive behaviour between populations at the simultaneously hermaphroditic snail *Lymnaea stagnalis*

**DOI:** 10.1101/2025.06.11.658905

**Authors:** Susanne Pekelharing, Yumi Nakadera, Takumi Saito, Joris M. Koene

## Abstract

Differences in reproductive behaviour can contribute to speciation by establishing reproductive isolation and can thus be a significant driving force in ecology and evolution. However, studies investigating intraspecific variation in reproductive behaviour remain scarce. In this study, we focused on a simultaneously hermaphroditic freshwater snail, *Lymnaea stagnalis*, a model organism whose reproductive behaviour is well documented, to examine behavioural differences among populations. In addition to a laboratory population, ten wild populations were sampled and standardised under laboratory conditions. Mating behaviour and egg production were quantified through observation experiments. The observed traits were analysed using a hierarchical Bayesian model to estimate the effects of each population on reproductive traits. Moreover, genetic information of the populations was estimated using low-coverage whole genome sequencing data, and phylogenetic signals for each trait were assessed. Our results revealed that populations exhibited distinct behavioural traits, and most prominent were the differences in male behaviours and the motivation to mate in the male role (after short and long sexual isolation). Egg production also clearly differed between populations. However, no phylogenetic signals were detected for any of the traits, suggesting that the diversification of behavioural traits among populations may result from evolutionary processes such as divergent selection or founder effects. This study highlights that reproductive behaviour can be highly diversified within a single species, and that assuming homogeneity in behaviour should be done with care.

## Introduction

Understanding the mechanisms of speciation is a central issue in evolution and ecology, and reproductive isolation is a central process of speciation (Coyne and Orr 2004). Significant diversity among closely related species in reproductive traits, including behaviour, is observed across many taxa (Pillay and Rymer 2012; Wilkins et al. 2013; Meiri et al. 2020), and these traits may have rapidly evolved as a result of both positive and relaxed selection (Svensson and Gosden 2007; Dapper and Wade 2020; Wiberg et al. 2021; Flacchi et al. 2024). Moreover, intraspecific divergence of reproductive traits can consequently lead to speciation via reproductive isolation (Coyne and Orr 2004; Hayashi et al. 2007; Maan and Seehausen 2011; Lipshutz et al. 2016). The basis for this sequential process should naturally be the diversity of reproductive traits within a species, and indeed significant variation has been observed in several animal taxa (e.g., Saarikettu et al. 2005; Ishikawa et al. 2006; Polihronakis 2006; Amézquita et al. 2009; González et al. 2013; Showalter et al. 2013; Goenaga et al. 2015; Bollatti et al. 2017; Olivero et al. 2017; Nakadera et al. 2020; Patlar et al. 2021). However, quantitative knowledge of consistent differences in reproductive traits, especially for behavioural differences between populations of a species, is still lacking for most taxa, although these may play a key role in the initial steps of reproductive isolation (Pillay and Rymer 2012).

One major hinderance for documenting evolutionary divergence in reproductive traits, that potentially lead to speciation, is phenotypic plasticity. Because, although the evolutionary diversity of reproductive traits within species can arise through various mechanisms, such as natural selection and founder effects (Matute 2013; Meiri et al. 2020), these differences can also result from phenotypic plasticity (Ford and Seigel 1989; Luo et al. 2010). In particular, the plasticity of reproductive behaviour has garnered increasing research attention in recent years (Dore et al. 2018; Iossa et al. 2019; Kirkpatrick and Sheldon 2022), with evidence suggesting that behavioural plasticity can affect several evolutionary processes (Caspi et al. 2022). To distinguish evolutionary differences from plastic, non-genetic changes, organisms under investigation need to be compared under standardised conditions to minimise the effects of plasticity.

A limited number of studies have examined intraspecific variation in reproductive behaviours under controlled conditions. One of the most extensively studied example can be the behavioural diversity in *Drosophila* species (e.g. Patty, 1975; Cohet and David 1980; Gromko and Newport, 1988; Saarikettu et al. 2005; Hurtado and Hasson, 2013; Singh and Singh, 2013). For several *Drosophila* species, laboratory stock populations originating from different geographic origins were primarily used to investigate variation in reproductive behaviours, such as the remating duration. These studies suggest that sexual selection operating at different stages of reproduction may drive behavioral divergence among populations. These laboratory stocks have typically been maintained under stable conditions for extended periods; therefore, the effects of plasticity can be largely controlled, while they may not be regarded equivalent to the wild populations. Similarly, variations in reproductive behaviours among laboratory strains of the house mouse (*Mus musculus* Linnaeus, 1758) are well documented (reviewed in Niel and Monks 2013; e.g. Levine, 1966; Batty 1978; Canastar and Maxson 2003). Another well-established model is the guppy (*Poecilia reticulata* Peters, 1859). Most research on guppies has focused on reproductive behaviours—particularly mate choice—among populations originating from habitats with differing levels of predation risk. These studies indicate that environmental differences can lead to divergence in reproductive behaviours at population level (reviewed in Brooks 2002; e.g. Godin and Briggs 1996; Houde and Hankes 1997; Grant et al. 2025).

In contrast, aside from research on such model organisms, most studies investigating intraspecific variation in reproductive behaviour rely on field observations or behavioural experiments of individuals collected from wild conditions (e.g. Ishikawa et al. 2006; González et al. 2013; Olivero et al. 2017). As a result, transgenerational and developmental plasticity cannot be adequately accounted for in such studies. In addition, all of the above-mentioned organisms are gonochoric species, and our knowledge of intraspecific variations in reproductive behaviours of hermaphrodites remains limited, although hermaphrodites may exhibit distinct evolutionary dynamics in reproductive traits compared to gonochoric species, due to the differences in sexual conflict mechanisms (Schärer et al. 2015). Therefore, to further understand the process underlying the diversification of reproductive traits, it is imperative to compare the reproductive behaviour of multiple populations under standardised conditions and to accumulate knowledge about the diversity of these traits.

Here, we focused on the great pond snail, *Lymnaea stagnalis* (Linnaeus, 1758) (Lymnaeidae, Gastropoda, Mollusca), which provides an excellent model system for investigating several biological disciplines, such as neurobiology, ecotoxicology, developmental biology and reproductive biology (Amorim et al. 2019; Fodor et al. 2020; Kuroda and Abe, 2020; Rivi et al., 2020; Koene et al. 2024). *Lymnaea stagnalis* is a simultaneously hermaphroditic species and especially within the contexts of reproductive and evolutionary biology this species can offer unique insights into the evolution of sexual conflict within an individual via sexual selection (Koene 2006). Consequently, substantial knowledge has been accumulated on the reproductive biology of *L. stagnalis* (e.g. Van Duivenboden and Ter Maat 1985; Koene et al. 2008; Hermann et al. 2009; Nakadera et al. 2014b).

The reproductive traits of *L. stagnalis*, including behaviours, are affected by several individual-level factors, such as body size, age, mating experience, and even learning (e.g. Koene et al. 2006, 2007, 2009, 2010; Koene and Ter Maat 2007; Hermann et al. 2009; Hoffer et al. 2012, 2017; Nakadera et al. 2014b, 2015; Moussaoui et al. 2018; Daupagne and Koene 2020; Swart et al. 2020; Álvarez et al. 2022). For instance, the body size positively influences egg productions (Koene et al. 2007; Nakadera et al. 2015), and the age has significant effects on their mating role preference, with younger individuals tending to mate in the male role (Hermann et al. 2009; Nakadera et al. 2015). Moreover, accessory gland proteins (ACPs), which are socially transferred from donors to recipients during insemination, trigger various physiological responses in recipients (Hakala et al. 2023). These responses include delayed egg laying (Koene et al. 2010; Nakadera et al. 2014b) and reduced sperm investment (in the next mating: Nakadera et al. 2014b), and induced behaviours that seem to be intended to avoid additional ejaculate receipt (Daupagne and Koene 2020). Such plasticity in reproductive traits is considered to be a consequence of sexual selection under complex sexual conflict in hermaphroditic species (Koene 2006).

However, despite its extensive use in experimental research, most studies on *L. stagnalis* have been limited to single populations, such as a laboratory breeding strain, and studies addressing the variation among populations in reproductive traits remain scarce (e.g. Puurtinen et al. 2007; Nakadera et al. 2020). This study aims to investigate the diversity of reproductive traits in *L. stagnalis* across populations by analysing mating behaviours and oviposition traits using standardised populations from the field using a Bayesian hierarchical modelling. With the findings we expect to reveal the mechanisms underlying the evolutionary diversification of reproductive traits and highlight the overlooked intraspecific variation in *L. stagnalis,* further enriching its role as a model organism.

## Materials and Methods

### Model organisms

*Lymnaea stagnalis* populations were collected from eleven localities across Europe (see location details in Table 1). These populations, except for the laboratory strain (LAB), have been cultured for six to seven generations at the breeding facility of the Vrije Universiteit Amsterdam, The Netherlands before the mating experiments. This cultivation period likely eliminated any possible influence of plasticity including transgenerational effects remaining from the field and provided standardisation for each population. The LAB population was kept in cohorts of the same age in large tanks (50 L) under standard conditions: laminar flow low-copper water at 20 ± 1°C, a 12:12 hour light-dark cycle, and water flow of 200 L/h. Other populations were housed in separate smaller tanks (3 L, at roughly the same density as the Lab population) and each population cohort was age synchronised (like the LAB population). These tanks had thin slits on two opposing sides to allow for water exchange when placed within a larger tank (170 L) under the same standard condition as indicted above. For all snails, a sufficient supply of butterhead lettuce (*Lactuca sativa*) was provided as food. The snails used for the experiments were randomly selected from individuals over 20 mm or larger and also all snails used were over 3-month-old and were all fully mature (Van Duivenboden 1983; Van Duivenboden and Ter Maat, 1985; Koene et al. 2008). Then, the snails were isolated in 625 mL perforated containers placed in a laminar flow tank under the same conditions, this means that the containers were filled with a volume of 460 ml. The donor snails were isolated for 7 days to increase their motivation to mate in the male role, as their prostate gland increases in size during this time which provides a permissive signal to mate in that role (De Boer et al. 1997; Koene and Ter Maat 2007). The recipient snails were isolated for 3-4 days to mitigate the effects of prior inseminations without increasing their motivation to mate in the male role (Koene et al. 2009, 2010; Nakadera et al. 2014a). All snails that died during the quarantine period were excluded from subsequent experiments.

**Table 1.** Information of *Lymnaea stagnalis* populations used in this study.

| Code | Locality | N | E | N (pairs) | Remark |
| --- | --- | --- | --- | --- | --- |
| AU | Salzburg, Austria | 47.799 | 13.391 | 19 |  |
| BE | Antwerp, Belgium | 51.031 | 4.216 | 19 |  |
| FI | Jyväskylä, Finland | 62.617 | 26.35 | 5 |  |
| FL | Lyon, France | 45.773 | 5.467 | 20 | The same as F2 in Nakadera et al. (2020) |
| FR | Rennes, France | 48.113 | -1.71 | 20 | The same as F1 in Nakadera et al. (2020) |
| GR | Thessaloniki, Greece | 41.25 | 23.21 | 19 |  |
| LAB | Eemnes, The Netherlands | 52.252 | 5.287 | 20 | Laboratory culture for over 50 years |
| M1 | Maastricht, The Netherlands | 50.822 | 5.676 | 17 |  |
| M2 | Maastricht, The Netherlands | 50.818 | 5.679 | 15 |  |
| UT | Utrecht, The Netherlands | 52.085 | 5.126 | 19 | The same as NL in Nakadera et al. (2020) |
| SW | Zürich, Switzerland | 47.475 | 8.729 | 13 |  |

### Experimental design

After the isolation period, the shell height (SH) of each snail - reliable predictor of body size (Koene et al. 2007; Koene et al. 2008) - was measured using a digital calliper (Traceable® Digital Calliper 8 in., VWR International, PA), and the snails were manually paired according to similarity in size to mitigate potential influences of size differences (Nakadera et al. 2015). In addition, all pairs used individuals of the same age to eliminate the effect of age differences (Hermann et al. 2009; Nakadera et al. 2015). In total, 186 pairs were set up for the behavioural experiment (Table 1). The donor snails were marked using a small dot of nail polish to distinguish them from the recipient snails. Subsequently, they were transferred to 625 mL containers without perforations, filled with approximately 460 mL water from the same tank in which they were isolated, and observed in the same room where the temperature was the same as the water temperature, 20 ± 1 °C. After the preparation, the 6-hour observation period was started, and the behaviour of the snail pairs was closely observed and recorded every 5 minutes (scan sampling; e.g. Palmeira et al. 2023). In this study, 16 experimental runs were conducted over several weeks because it is impossible to observe all pairs within a single observation moment. The observations took place once or twice a week and involved 2 to 3 populations, each with approximately 5 pairs of snails. To reduce the potential influence of day-to-day variations on mating behaviour, the used populations were randomly selected and the observation days were varied. Both mating behaviours as a male and as a female were quantified according to previous studies (Van Duivenboden and Ter Maat 1988; Moussaoui et al. 2018; Álvarez et al. 2022; Palmeira et al. 2023; Table 2). Additionally, we noted whether the potential sperm donor was positioned on top of the potential sperm receiver during some female behaviours, and recorded the frequency of biting per snail as in Moussaoui et al. (2018). After the observation period, the recipient snails were isolated again for 7 days in clean perforated containers in the large flowthrough tank to allow for egg laying. All the egg masses were collected three times during the 7 days (always including the 7^th^ day). As a results, we collected 624 egg masses from the 179 recipients and this data was used for subsequent analyses. To quantify the egg production, we scanned egg masses using a digital scanner (CanoScan LiDE 220). Then, individual egg capsule numbers within each egg mass (hereafter, egg number) were counted and the area of five egg capsules was quantified (hereafter egg size) that were randomly selected from each egg mass were measured on the digital scan using Fiji (Schindelin et al. 2012).

**Table 2.**
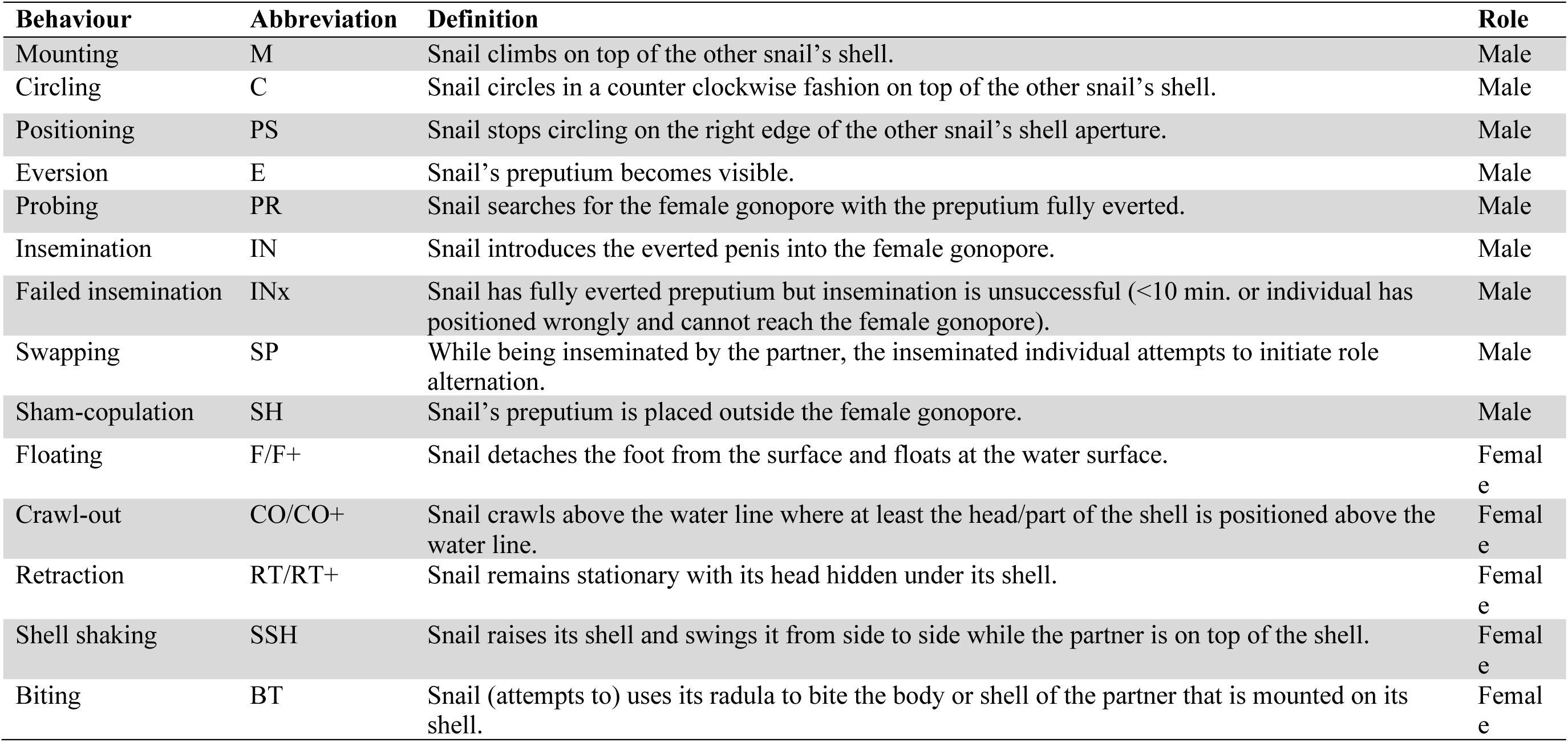
Definitions of the recorded behaviours, their abbreviations and the sexual role in which they are performed. + mark means the potential sperm donor was positioned on top of the potential sperm receiver

| Behaviour | Abbreviation | Definition | Role |
| --- | --- | --- | --- |
| Mounting | M | Snail climbs on top of the other snail's shell. | Male |
| Circling | C | Snail circles in a counter clockwise fashion on top of the other snail's shell. | Male |
| Positioning | PS | Snail stops circling on the right edge of the other snail's shell aperture. | Male |
| Eversion | E | Snail's preputium becomes visible. | Male |
| Probing | PR | Snail searches for the female gonopore with the preputium fully everted. | Male |
| Insemination | IN | Snail introduces the everted penis into the female gonopore. | Male |
| Failed insemination | INx | Snail has fully everted preputium but insemination is unsuccessful (<10 min. or individual has positioned wrongly and cannot reach the female gonopore). | Male |
| Swapping | SP | While being inseminated by the partner, the inseminated individual attempts to initiate role alternation. | Male |
| Sham-copulation | SH | Snail's preputium is placed outside the female gonopore. | Male |
| Floating | F/F+ | Snail detaches the foot from the surface and floats at the water surface. | Female |
| Crawl-out | CO/CO+ | Snail crawls above the water line where at least the head/part of the shell is positioned above the water line. | Female |
| Retraction | RT/RT+ | Snail remains stationary with its head hidden under its shell. | Female |
| Shell shaking | SSH | Snail raises its shell and swings it from side to side while the partner is on top of the shell. | Female |
| Biting | BT | Snail (attempts to) uses its radula to bite the body or shell of the partner that is mounted on its shell. | Female |

### Genetic analysis

To examine the genetic relationship between the populations, we utilised low-coverage whole genome sequencing (lcWGS). The lcWGS is a cost-effective way to obtain data from whole genomes (Lou et al. 2021) and can eliminate the effect on genetic relationship of evolutionary artifact such as incomplete lineage sorting, introgressive hybridization, and mito-nuclear discordance which are often observed in freshwater gastropods (Hirano et al. 2019; David et al. 2022; Stelbrink et al. 2024). The lcWGS were conducted using field caught individuals of each population (except for LAB). Then, we collected foot tissues after anesthetisation with 50 mM MgCl2 for DNA extraction. We snap froze these samples in liquid nitrogen and stored them at -20 C. We extracted total DNA using DNeasy Blood and Tissue Kit (Qiagen, Hilden, Germany). The PCR-free library preparation and Illumina (CA, USA) sequencing (PE150) was conducted by Novogene (Cambridge, UK).

From all the data, two individuals after de-multiplexing from each population were used in the subsequent analysis. The adaptor and low-quality (< Q20) sequences were removed using Atria v3.2.2-1-Linux (Chuan et al. 2021) and paired sequences were merged with a possible 20bp extension using BBMerge-auto.sh in BBTools (Bushnell et al. 2017). Then, genomic distances were estimated using Skmer (Sarmashghi et al. 2019) which is specialised for lcWGS and based on the estimated distances under JC69 model (Jukes and Cantor 1969), and the average distance of two individuals were calculated for each population. Then, a taxon addition phylogeny with the minimum-evolution principle (Desper and Gascuel 2002) was estimated using FastME 2.0 (Lefort et al. 2015).

### Statistical modelling

Based on previous studies (Hermann et al. 2009; Nakadera et al. 2015), body size and age are influential factors on the reproductive traits of *L. stagnalis*. To comprehensively model these factors and to elucidate differences in reproductive traits across populations, we employed a hierarchical Bayesian modelling. All analyses were performed using *rstan* package under R version 4.2.2 (Carpenter et al. 2017; R Core Team 2022), and results were visualised using *ggplot2*, *ggridges* and *ggmcmc* packages (Fernández-i-Marín 2016; Wickham 2016; Wilke 2024).

Frequency of individual behaviours were separately recorded as the sperm donor’s male (donor male) and female (donor female) behaviours, and the sperm recipient’s female (recipient female) and male (recipient male) behaviours. These data were summarised thorough principal component analysis (PCA) with standardisation using *prcomp* function in R. Each of PC1 values (i.e., donor/male, donor/female, recipient/male and recipient/female) was modelled with a normal distribution under the hierarchical Bayesian framework and the size and age were incorporated into the models as each dependent variable (Eqs. 1–2). SH was used for the indicator for the body size and the number of days after egg masses collecting was employed as the indicator of the age. Effect of each population were included as a fixed effect *Pop*_*g*_ for each population, because population effects cannot be assumed to follow a linear trend across groups. In addition, uncontrolled differences among 16 experimental runs were incorporated as a random effect ε_*ex*_. The body size was usually correlated with the age in *L. stagnalis* (Koene et al. 2008; Hermann et al. 2009; Nakadera et al. 2015); however, the correlation and variance inflation factor (VIF) for a linear regression (Eq. 2) were modest in our dataset (Donor: *r* = 0.64, VIF = 1.73; Recipient: *r* = 0.71, VIF = 2.05). Thus, to address potential multicollinearity, the intercept (β_*pc*1_) and coefficients (β_*pc*2_ and β_*pc*3_) were assigned normal distributions with a mean of 0 and a variance of 10^2^ as weakly informative prior distributions. For ε_*ex*_ and *Pop*_*g*_, we set normal distributions, with mean of 0 and variance of σ_*ex*_ and σ_*pop*_), respectively. Variances (σ_*pc*_, σ_*ex*_, and σ_*pop*_) were assigned half-Cauchy distributions with a scale parameter (γ) of 4 as weak-informative prior distribution. The size and age data were standardized (mean = 0 and variance = 1) prior to Markov chain Monte Carlo (MCMC) sampling. Then, MCMC runs with default settings were performed for 10×10^3^ iterations across four parallel chains, sampling every fifth iterations after a burn-in of 5×10^3^ iterations. The convergence of the MCMC runs were assessed using the effective samples size (ESS; > 200), the potential scale reduction statistic (*Rhat* < 1.02; Gelman and Rubin 1992), and inspection of the trace plots. Estimated effects were considered statistically significant when probability of direction (PD: Makowski et al. 2019) exceeded 97.5% on the posterior distribution.

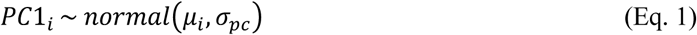

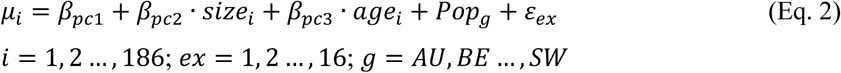

The mating success was assessed independently for the donor and recipient as a primary representative of mating behaviours. Mating success was defined as insemination initiated within 6 hours considered successful (1) and failure (0) otherwise. A hierarchical Bayesian model with a Bernoulli distribution was constructed, applying binary logistic regression (Eqs. 3–4). As with the PC1 model, the body size and age were integrated into the model as predictors (Eq. 4). For all the coefficients, we set normal distributions with mean of 0 and variance of 10^2^. For ε*_ex_* and *Pop_g_*, we set normal distributions with mean of 0 and variance of σ*_ex_* and σ*_pop_* For the variances (σ*_ex_* and σ*_pop_*), we set half-Cauchy distributions with γ of 4. The age and size data were standardized and MCMC runs, assessment of the convergence and visualization were performed in the same manner as the PC1 model.

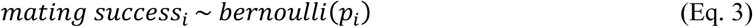

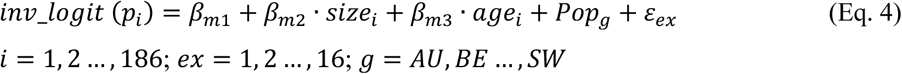

Total egg production over 7 days was expressed as the product of the egg capsule number and the average size of the egg capsules. In general, there may be a trade-off between egg number and size (e.g., Fleming and Gross 1990; Christians 2000; Charnov and Ernest 2006); however, in *L. stagnalis*, an existence of the trade-off remains unclear. On the other hand, the egg number decreased and the egg size increased following mating (Hoffer et al. 2012; Swart et al. 2020). Egg number and size were therefore modelled separately to avoid an overly complex model, since the primary focus of this model is variation on egg laying among populations.

The egg size was modelled with a normal distribution, incorporating the body size (SH), age, mating status as a recipient (i.e., inseminated by the donor), and the total egg number as predictors (Eqs. 5–6). Previous studies demonstrated a certain impact on egg production for the former three variables (Koene et al. 2006, 2009, 2010; Hoffer et al. 2012; Nakadera et al., 2014b; Swart et al. 2020) and the latter factor can account for potential the trade-off relationship between the egg number and size. A corelation between the age and size was moderate (*r* = 0.71, VIF_size = 1.38, VIF_age = 1.27). All coefficients were assigned normal distributions with mean of 0 and variance of 10^2^, and ε*_ex_* and *Pop_g_* followed normal distributions with mean of 0 and variance of σ*_ex_* and σ*_pop_*. For the variances (σ*_eggav_*, σ*_ex_* and σ*_pop_*), we set half-Cauchy distributions with γ of 4. Standardization and MCMC protocols were consistent with previous models.

Egg number was modelled with a negative binomial distribution and exponential regression (Eqs. 7–8). The size, age, mating, and the average egg size were integrated into the model as predictors (correlation between the size and age: VIF_size = 1.15, VIF_age = 1.16). Priors and MCMC procedures paralleled those employed in the egg size model.

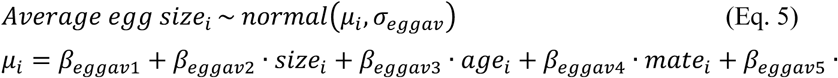

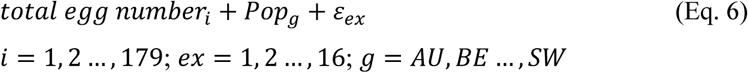

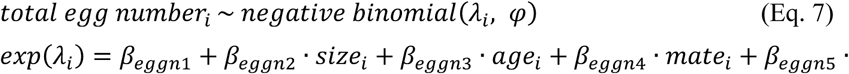

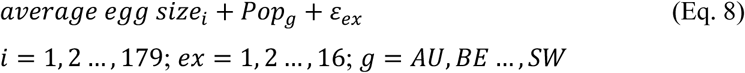

### Test for phylogenetic signals

Finally, we tested phylogenetic signals to examine whether reproductive traits were associated with phylogenetic relationships under a Brownian motion (BM) model. Phylogeny obtained from lcWGS (Figure S1) were compared with the mean PC1 values, mating success rates, total egg numbers and average egg sizes across populations. Because each phylogenetic signal index provides different properties (Münkemüller et al. 2012), we computed and statistically tested four representative indices: Abouheif’s *C*mean (Abouheif 1999), Moran’s *I* (Gittleman and Kot 1990), and Blomberg’s *K* (Blomberg et al. 2003) using *phylosignal* package in R version 4.2.2 (Keck et al. 2016).

## Results

### Behavioural analyses

Not all matings were performed in the intended order, which would have been 7d isolated individuals performing the male role first and 4d isolated animals performing the female role first. When the intended donor performed the male behaviour first, PC1 accounted for 28.4% of the variance, while PC2 explained 13.4% (Table S1). The average PC1 values for each population ranged from -0.59 (UT) to 0.91 (LAB) (Figs 1–2, Table S2). When the intended donor performed the female behaviour first, PC1 explained 21.0% of the variance, and PC2 accounted for 14.4%, with average PC1 values ranging from -1.00 (M2) to 0.96 (BE). When the intended recipient performing the female behaviour first, PC1 accounted for 19.4% of the variance, and PC2 explained 15.7%, with average PC1 values ranging from -0.49 (M2) to 0.52 (LAB), while PC1 for the intended recipient performing the male behaviour first accounted for 34.4% of the variance, and PC2 explained 16.6%, with average PC1 values ranging from -1.25 (M2) to 2.36 (BE).

**Figure 1.**
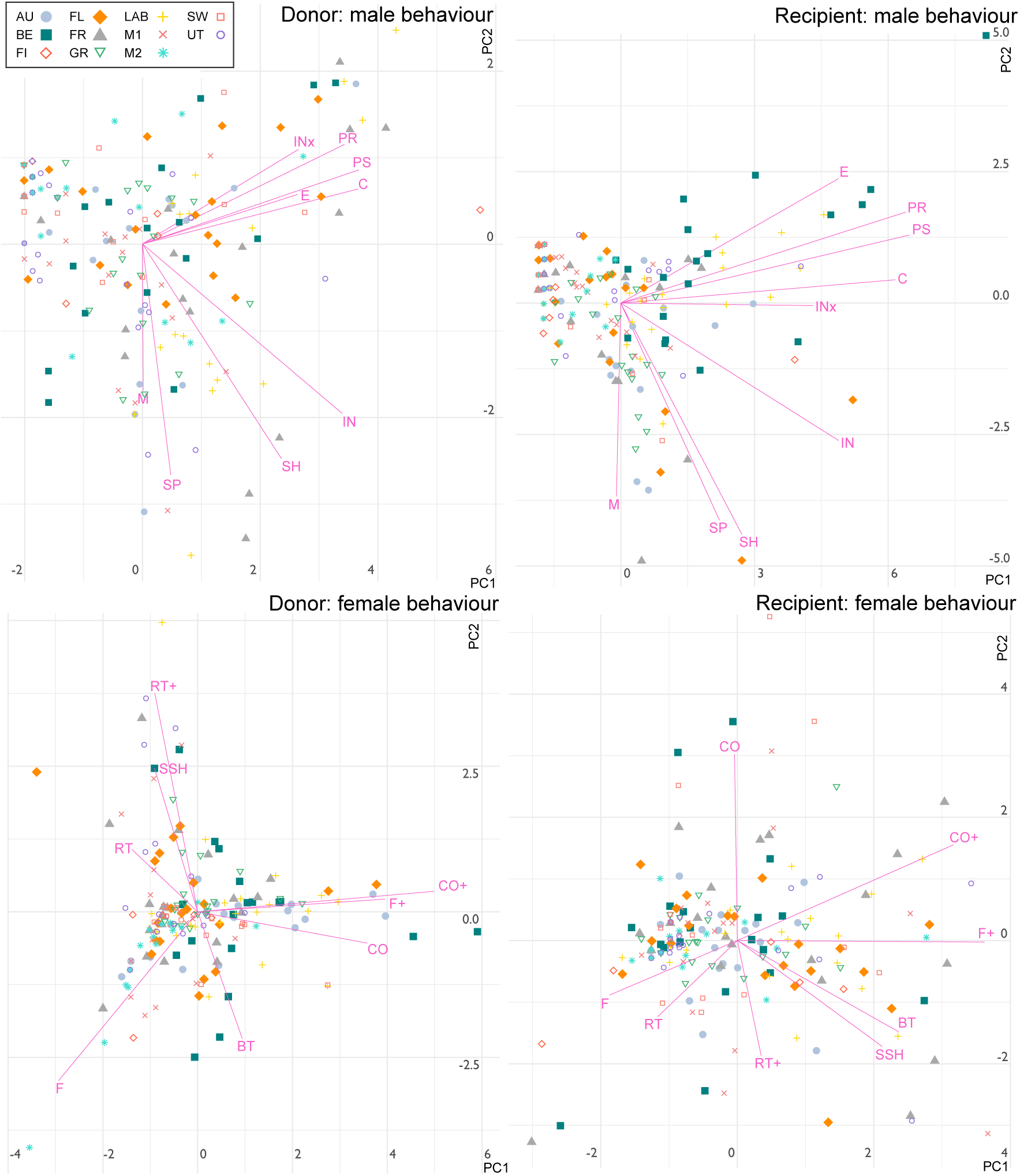
Principal component plots for behavioural observation of intended Donors and Intended recipients during six hours. Symbols indicate populations (Table 1) and lines on graphs displays loadings for each behaviour (Table 2). See also Dataset A1 for full results.

**Figure 2.**
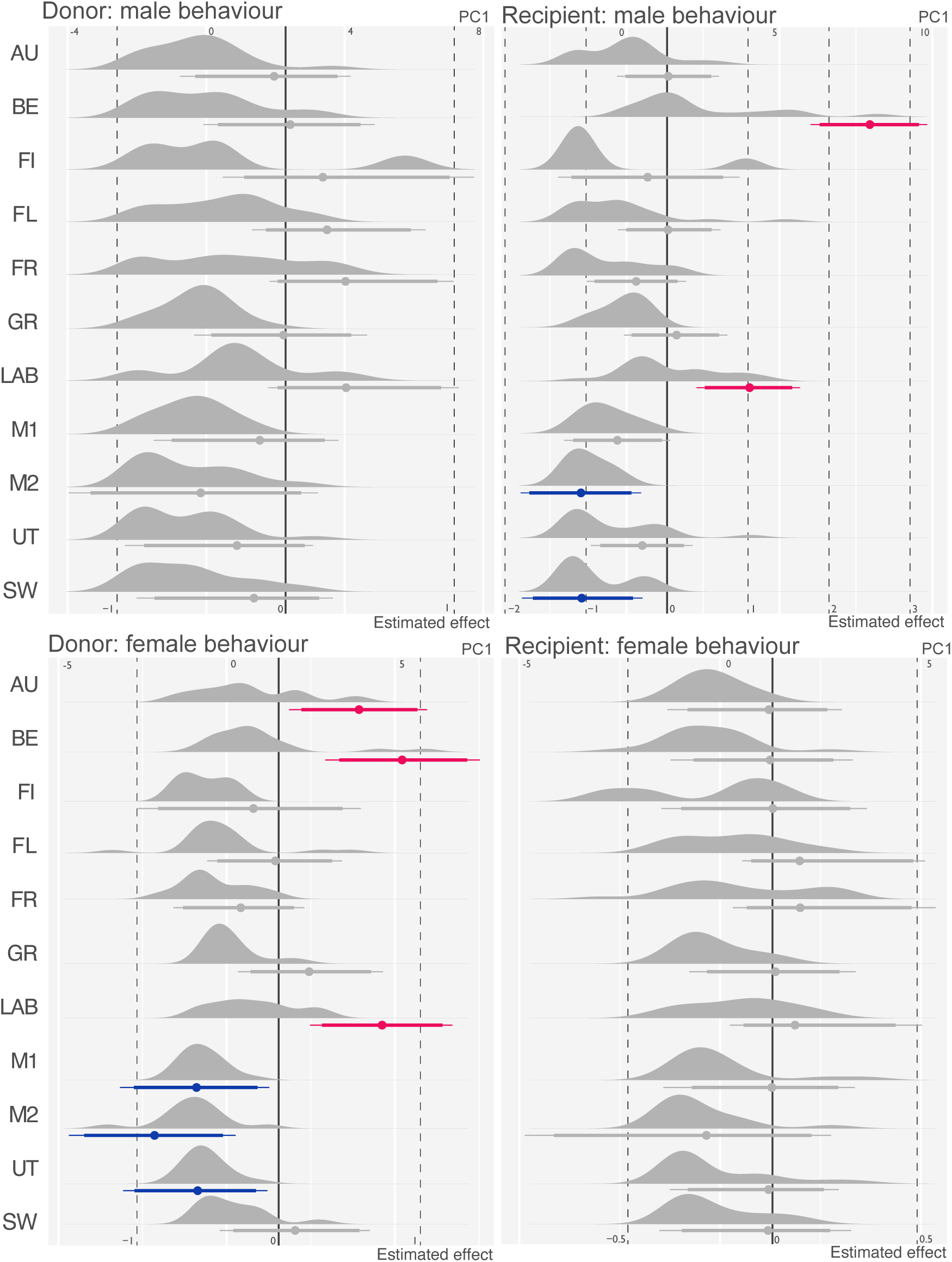
Distributions of principal component 1 (PC1) for each population and estimated effects on PC1s from each population. Codes indicate populations (Table 1). Upper axis displays value for PC1s and bottom axis displays value for effects. Effects are presented in Bayesian confidence interval (CI). Dots on CI indicate mean values. Bold lines indicate 90% CI and thin lines indicate 95% CI. Negative significant effects are shown in blue, and positive significant effects shown in pink. See also Table 3 and Dataset A1 for full results.

**Table 3.**
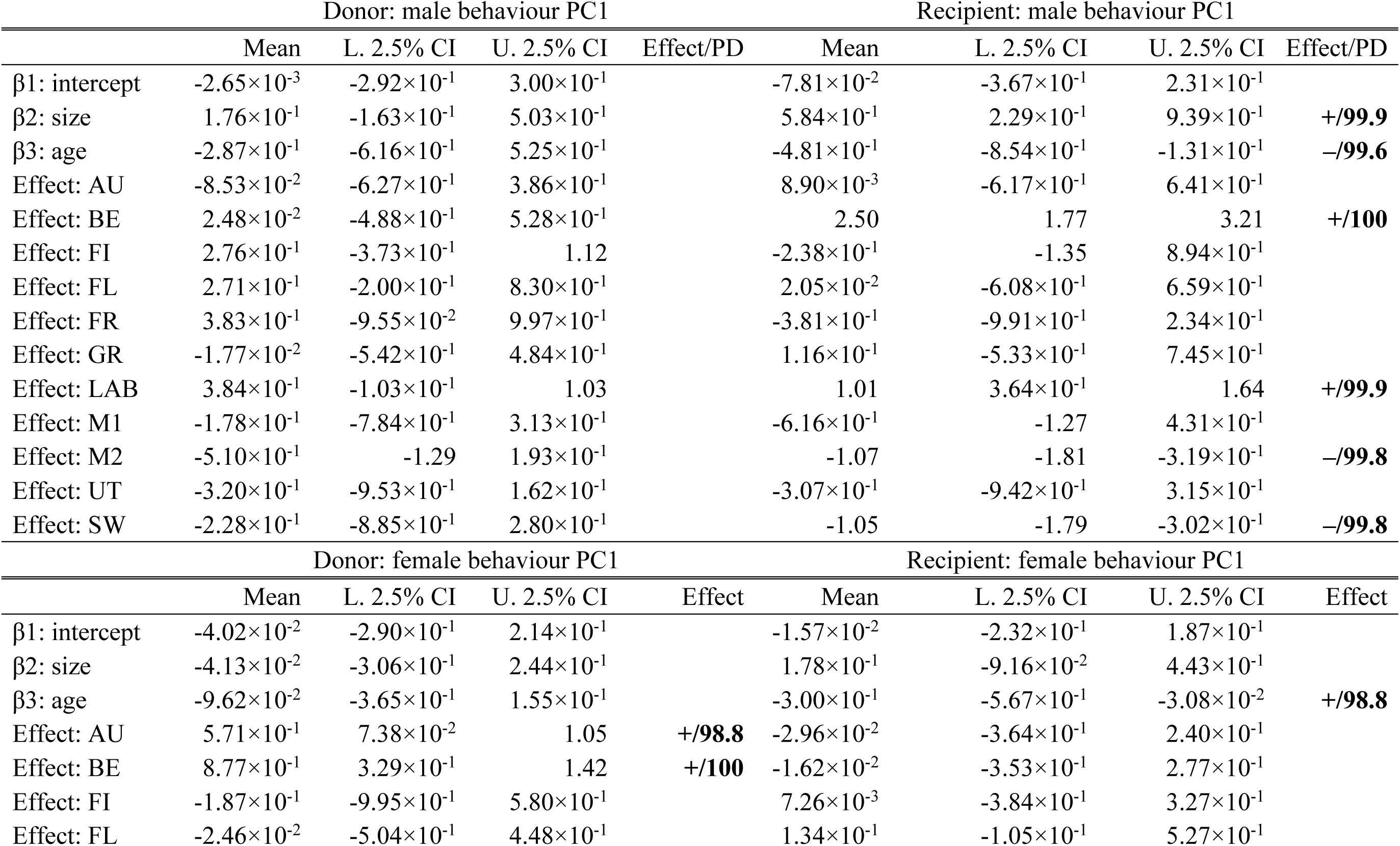
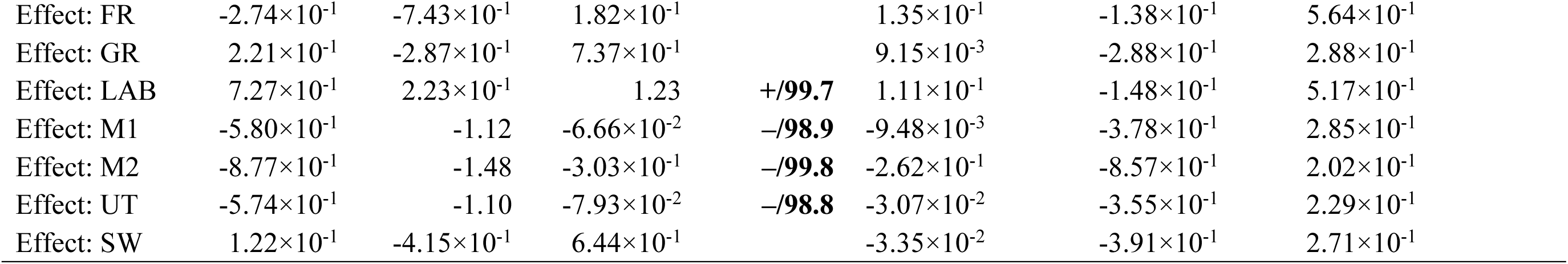
Estimates from modelling for reproductive behaviours.

|  | Donor: male behaviour PC1 |  |  |  | Recipient: male behaviour PC1 |  |  |  |
| --- | --- | --- | --- | --- | --- | --- | --- | --- |
|  | Mean | L. 2.5% CI | U. 2.5% CI | Effect/PD | Mean | L. 2.5% CI | U. 2.5% CI | Effect/PD |
| $\beta 1$: intercept | $-2.65 \times 10^{-3}$ | $-2.92 \times 10^{-1}$ | $3.00 \times 10^{-1}$ | | $-7.81 \times 10^{-2}$ | $-3.67 \times 10^{-1}$ | $2.31 \times 10^{-1}$ | |
| $\beta 2$: size | $1.76 \times 10^{-1}$ | $-1.63 \times 10^{-1}$ | $5.03 \times 10^{-1}$ | | $5.84 \times 10^{-1}$ | $2.29 \times 10^{-1}$ | $9.39 \times 10^{-1}$ | **+/99.9** |
| $\beta 3$: age | $-2.87 \times 10^{-1}$ | $-6.16 \times 10^{-1}$ | $5.25 \times 10^{-1}$ | | $-4.81 \times 10^{-1}$ | $-8.54 \times 10^{-1}$ | $-1.31 \times 10^{-1}$ | **-/99.6** |
| Effect: AU | $-8.53 \times 10^{-2}$ | $-6.27 \times 10^{-1}$ | $3.86 \times 10^{-1}$ | | $8.90 \times 10^{-3}$ | $-6.17 \times 10^{-1}$ | $6.41 \times 10^{-1}$ | |
| Effect: BE | $2.48 \times 10^{-2}$ | $-4.88 \times 10^{-1}$ | $5.28 \times 10^{-1}$ | | 2.50 | 1.77 | 3.21 | **+/100** |
| Effect: FI | $2.76 \times 10^{-1}$ | $-3.73 \times 10^{-1}$ | 1.12 | | $-2.38 \times 10^{-1}$ | -1.35 | $8.94 \times 10^{-1}$ | |
| Effect: FL | $2.71 \times 10^{-1}$ | $-2.00 \times 10^{-1}$ | $8.30 \times 10^{-1}$ | | $2.05 \times 10^{-2}$ | $-6.08 \times 10^{-1}$ | $6.59 \times 10^{-1}$ | |
| Effect: FR | $3.83 \times 10^{-1}$ | $-9.55 \times 10^{-2}$ | $9.97 \times 10^{-1}$ | | $-3.81 \times 10^{-1}$ | $-9.91 \times 10^{-1}$ | $2.34 \times 10^{-1}$ | |
| Effect: GR | $-1.77 \times 10^{-2}$ | $-5.42 \times 10^{-1}$ | $4.84 \times 10^{-1}$ | | $1.16 \times 10^{-1}$ | $-5.33 \times 10^{-1}$ | $7.45 \times 10^{-1}$ | |
| Effect: LAB | $3.84 \times 10^{-1}$ | $-1.03 \times 10^{-1}$ | 1.03 | | 1.01 | $3.64 \times 10^{-1}$ | 1.64 | **+/99.9** |
| Effect: M1 | $-1.78 \times 10^{-1}$ | $-7.84 \times 10^{-1}$ | $3.13 \times 10^{-1}$ | | $-6.16 \times 10^{-1}$ | -1.27 | $4.31 \times 10^{-1}$ | |
| Effect: M2 | $-5.10 \times 10^{-1}$ | -1.29 | $1.93 \times 10^{-1}$ | | -1.07 | -1.81 | $-3.19 \times 10^{-1}$ | **-/99.8** |
| Effect: UT | $-3.20 \times 10^{-1}$ | $-9.53 \times 10^{-1}$ | $1.62 \times 10^{-1}$ | | $-3.07 \times 10^{-1}$ | $-9.42 \times 10^{-1}$ | $3.15 \times 10^{-1}$ | |
| Effect: SW | $-2.28 \times 10^{-1}$ | $-8.85 \times 10^{-1}$ | $2.80 \times 10^{-1}$ | | -1.05 | -1.79 | $-3.02 \times 10^{-1}$ | **-/99.8** |
|  | Donor: female behaviour PC1 |  |  |  | Recipient: female behaviour PC1 |  |  |  |
|  | Mean | L. 2.5% CI | U. 2.5% CI | Effect | Mean | L. 2.5% CI | U. 2.5% CI | Effect |
| $\beta 1$: intercept | $-4.02 \times 10^{-2}$ | $-2.90 \times 10^{-1}$ | $2.14 \times 10^{-1}$ | | $-1.57 \times 10^{-2}$ | $-2.32 \times 10^{-1}$ | $1.87 \times 10^{-1}$ | |
| $\beta 2$: size | $-4.13 \times 10^{-2}$ | $-3.06 \times 10^{-1}$ | $2.44 \times 10^{-1}$ | | $1.78 \times 10^{-1}$ | $-9.16 \times 10^{-2}$ | $4.43 \times 10^{-1}$ | |
| $\beta 3$: age | $-9.62 \times 10^{-2}$ | $-3.65 \times 10^{-1}$ | $1.55 \times 10^{-1}$ | | $-3.00 \times 10^{-1}$ | $-5.67 \times 10^{-1}$ | $-3.08 \times 10^{-2}$ | **+/98.8** |
| Effect: AU | $5.71 \times 10^{-1}$ | $7.38 \times 10^{-2}$ | 1.05 | **+/98.8** | $-2.96 \times 10^{-2}$ | $-3.64 \times 10^{-1}$ | $2.40 \times 10^{-1}$ | |
| Effect: BE | $8.77 \times 10^{-1}$ | $3.29 \times 10^{-1}$ | 1.42 | **+/100** | $-1.62 \times 10^{-2}$ | $-3.53 \times 10^{-1}$ | $2.77 \times 10^{-1}$ | |
| Effect: FI | $-1.87 \times 10^{-1}$ | $-9.95 \times 10^{-1}$ | $5.80 \times 10^{-1}$ | | $7.26 \times 10^{-3}$ | $-3.84 \times 10^{-1}$ | $3.27 \times 10^{-1}$ | |
| Effect: FL | $-2.46 \times 10^{-2}$ | $-5.04 \times 10^{-1}$ | $4.48 \times 10^{-1}$ | | $1.34 \times 10^{-1}$ | $-1.05 \times 10^{-1}$ | $5.27 \times 10^{-1}$ | |

|  |  |  |  |  |  |  |  |
| --- | --- | --- | --- | --- | --- | --- | --- |
| Effect: FR | $-2.74 \times 10^{-1}$ | $-7.43 \times 10^{-1}$ | $1.82 \times 10^{-1}$ | | $1.35 \times 10^{-1}$ | $-1.38 \times 10^{-1}$ | $5.64 \times 10^{-1}$ |
| Effect: GR | $2.21 \times 10^{-1}$ | $-2.87 \times 10^{-1}$ | $7.37 \times 10^{-1}$ | | $9.15 \times 10^{-3}$ | $-2.88 \times 10^{-1}$ | $2.88 \times 10^{-1}$ |
| Effect: LAB | $7.27 \times 10^{-1}$ | $2.23 \times 10^{-1}$ | 1.23 | **+/99.7** | $1.11 \times 10^{-1}$ | $-1.48 \times 10^{-1}$ | $5.17 \times 10^{-1}$ |
| Effect: M1 | $-5.80 \times 10^{-1}$ | -1.12 | $-6.66 \times 10^{-2}$ | **-/98.9** | $-9.48 \times 10^{-3}$ | $-3.78 \times 10^{-1}$ | $2.85 \times 10^{-1}$ |
| Effect: M2 | $-8.77 \times 10^{-1}$ | -1.48 | $-3.03 \times 10^{-1}$ | **-/99.8** | $-2.62 \times 10^{-1}$ | $-8.57 \times 10^{-1}$ | $2.02 \times 10^{-1}$ |
| Effect: UT | $-5.74 \times 10^{-1}$ | -1.10 | $-7.93 \times 10^{-2}$ | **-/98.8** | $-3.07 \times 10^{-2}$ | $-3.55 \times 10^{-1}$ | $2.29 \times 10^{-1}$ |
| Effect: SW | $1.22 \times 10^{-1}$ | $-4.15 \times 10^{-1}$ | $6.44 \times 10^{-1}$ | | $-3.35 \times 10^{-2}$ | $-3.91 \times 10^{-1}$ | $2.71 \times 10^{-1}$ |

Based on the PCA results for the intended donor’s male behaviour (Fig. 1), PC1 was primarily associated with eversion (E), probing (PR), positioning (PS), circling (C) and failed insemination (Inx), while PC2 was mainly associated with mounting (M), swapping (SP) and sham-copulation (SH). Insemination (IN) represented an intermediate behaviour between these two axes. For the intended donor’s female behaviour (Fig. 1), crawl-out (CO/CO+) and floating (F+) were related with PC1, whereas retraction (RT+), shell shaking (SSH) and biting (BT) were linked to PC2, with the latter showing opposite direction. Retraction (RT) and floating (F) were identified as intermediate behaviours between these two axes, although each has a different direction. In the intended recipient’s female behaviour (Fig. 1), most behaviours exhibited intermediate directions between these two axes, while floating (F+) was associated with PC1 and crawl-out (CO) with PC2. In the intended recipient’s male behaviour, a similar pattern to the intended donor’s male behaviour can be seen in the PC1 and PC2 (Fig. 1).

Although PC values for each population considerably overlapped, distinct trends were observed (Fig. 2). Hierarchical models estimated no significant effects from each population on PC1 for intended donor male behaviour. However, for PC1 of intended recipient male behaviour, positive effects were estimated for the BE and LAB populations, while negative effects were estimated for M2 and SW (Fig. 2). Between the donor and recipient, the same populations exhibited relatively similar effects on male behaviour, although no population showed significant effects for both groups. For intended donor female behaviour (Fig. 2), positive effects were estimated for the AU, BE and LAB populations, while negative effects were estimated for M1, M2 and UT. No significant effects were estimated for intended recipient female behaviour (Fig. 2). For PC2s, no significant effects were estimated across all populations (results not shown). In the models for PC1s, the age exhibited a significant positive dependence with the size (Table 3). For male recipient behaviour, the size had a significant positive dependence and the age had a significant negative dependence with PC1 (Table 3). In addition, a positive dependence of the age with PC1 was estimated for recipient female behaviour, (Table 3).

### Mating success

The populations’ average mating success rate (i.e. successful insemination; Fig. 3, Table S2) for the donor ranged from 0.33 (SW) to 0.8 (LAB), while for the recipient the range was 0.20 (FI and M2) to 0.90 (LAB). The hierarchical model for the donor estimated a negative effect, meaning that the population had a significantly lower mating success than the other population, on the mating success for M2, positive effects (i.e., higher mating success) for the recipients of BE and GR and negative effects for M2 and SW. The donor and recipient populations generally displayed similar estimated effect directions (i.e. positive/negative), although opposite effects were observed for BE and FR. For example, BE showed a positive effect for recipients but not for donors, with credible intervals (CIs) including zero (compare left and right in Fig. 3). The age exhibited a significant negative dependence with the mating success in both the donor and recipients, suggesting that older individuals have less mating success in general (Table 4).

**Figure 3.**
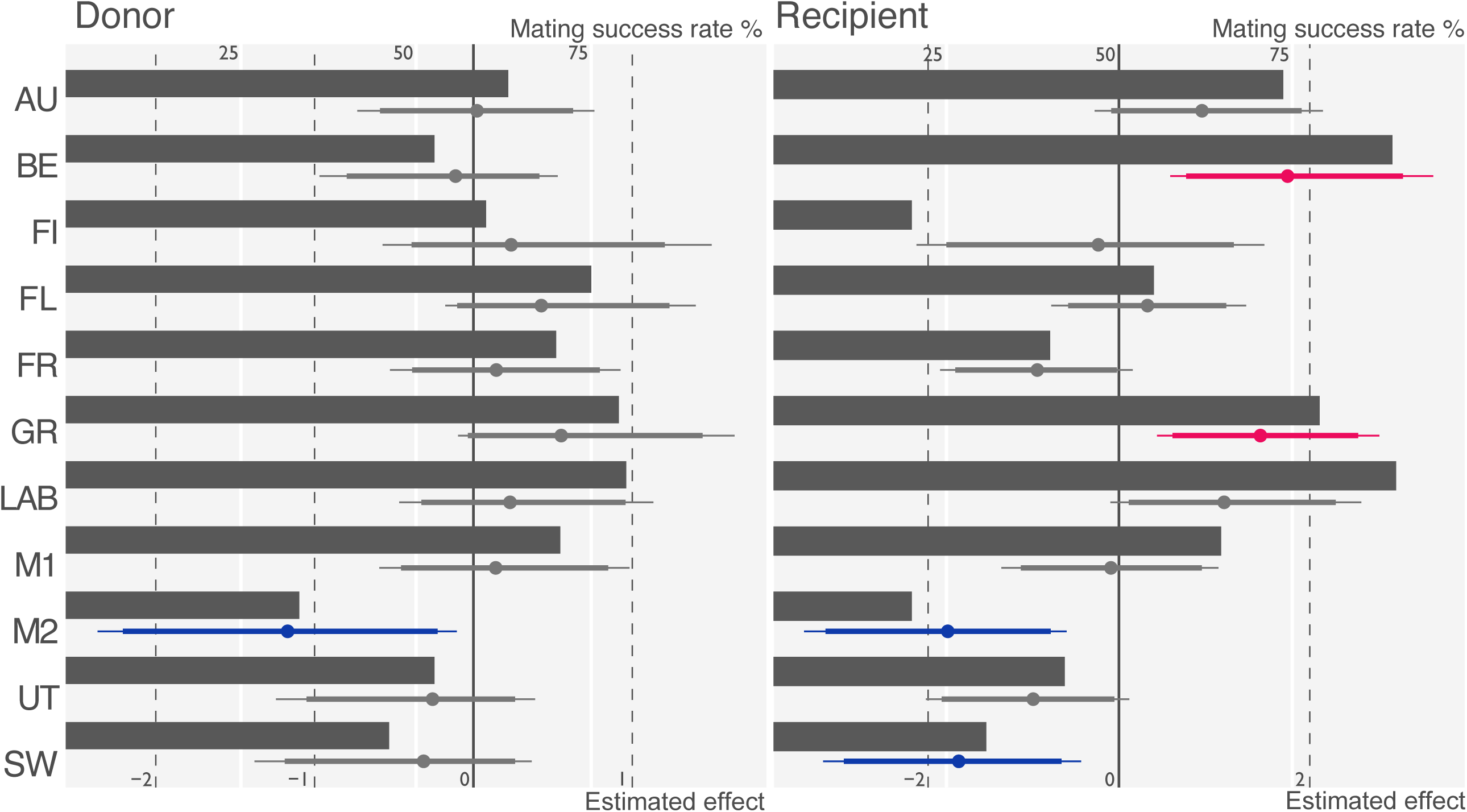
Male mating success rate and estimated effects on success rate from each population. Codes indicate populations (Table 1). Upper axis displays rate and bottom axis displays value for effects. Effects are presented as Bayesian confidence interval (CI). Dots on CI indicate mean value. Bold lines indicate 90% CI and thin lines indicate 95% CI. Negative significant effects are shown in blue, and positive significant effects shown in pink. See also Table 4 and Dataset A1 for full results.

**Table 4.** Estimates from modelling for mating success as male.

### Egg production

The average egg size (Fig. 4, Table S2) varied between populations from 1.057 (LAB) to 1.373 mm^2^ (FR), and the average egg number ranged from 138 (BE) to 283 (AU). The hierarchical model for the egg size estimated negative effects (i.e. a smaller egg size) for BE, LAB, M1, M2 and UT, whereas positive effects for FR, FL and SW. The model for the egg number only indicates a negative effect (i.e. a fewer number of eggs) for SW (Table 5). The model for the egg size indicates a significant negative dependence with egg number. Likewise, the model for the egg number estimated a significant negative dependence with egg size. The egg number model estimated a significant positive dependence on body size and a negative dependence on the age (Table 5).

**Figure 4.**
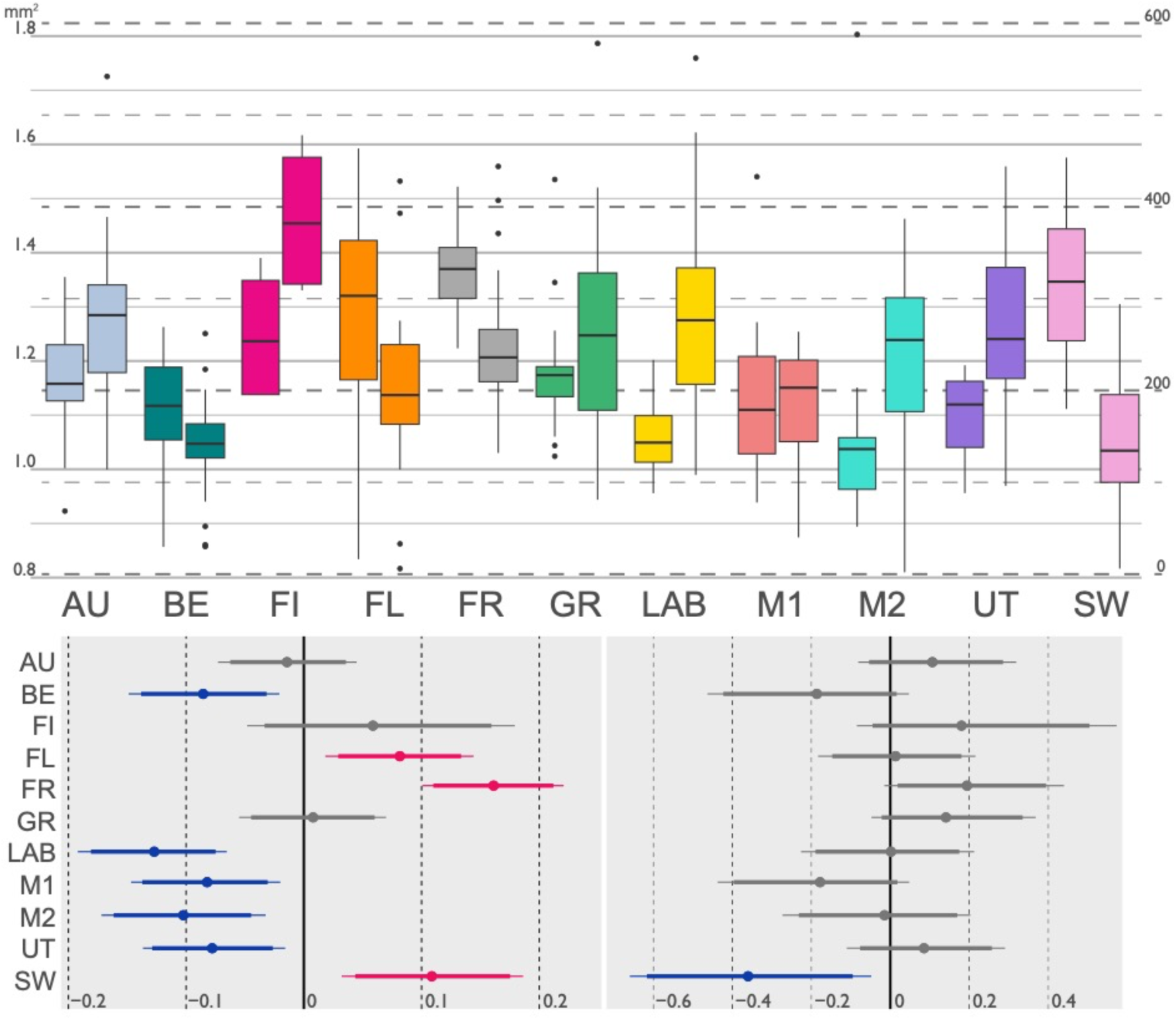
Box plots for egg size and number and estimated effects on egg size and number from each population. In the upper graph, left boxplot indicates average size of egg capsules (see left axis) and right boxplot indicates number of egg capsules (see count of right axis). Codes and colours indicate populations (Table 1). In the lower graph, left horizontal bar plot displays estimates of effects for egg size and right plot displays estimates of effects for egg number. Effects are presented in Bayesian confidence interval (CI). Dots on CI indicate mean value. Bold lines indicate 90% CI and thin lines indicate 95% CI. Negative significant effects are shown in blue, and positive significant effects shown in pink. See also Table 5 and Dataset A1 for full results.

**Table 5.** Estimates from modelling for egg size and egg number.

| Recipient: egg size |  |  |  |  |
| --- | --- | --- | --- | --- |
|  | Mean | Lower 2.5% CI | Upper 2.5% CI | Effect/PD |
| $\beta 1$: intercept | 1.19 | 1.15 | 1.23 | |
| $\beta 2$: size | $1.29 \times 10^{-2}$ | $-2.31 \times 10^{-2}$ | $4.89 \times 10^{-2}$ | |
| $\beta 3$: age | $-2.20 \times 10^{-2}$ | $-5.73 \times 10^{-2}$ | $1.43 \times 10^{-2}$ | |
| $\beta 4$: mate | $-9.39 \times 10^{-4}$ | $-4.49 \times 10^{-2}$ | $4.30 \times 10^{-2}$ | |
| $\beta 5$: egg no. | $-2.90 \times 10^{-2}$ | $-5.29 \times 10^{-2}$ | $-5.34 \times 10^{-3}$ | **−/99.3** |
| Effect: AU | $-7.05 \times 10^{-3}$ | $-6.66 \times 10^{-2}$ | $5.12 \times 10^{-2}$ | |
| Effect: BE | $-7.91 \times 10^{-2}$ | $-1.42 \times 10^{-1}$ | $-1.44 \times 10^{-2}$ | **−/99.2** |
| Effect: FI | $6.66 \times 10^{-2}$ | $-4.18 \times 10^{-2}$ | $1.86 \times 10^{-1}$ | |
| Effect: FL | $8.76 \times 10^{-2}$ | $2.45 \times 10^{-2}$ | $1.51 \times 10^{-1}$ | **+/99.6** |
| Effect: FR | $1.68 \times 10^{-1}$ | $1.07 \times 10^{-1}$ | $2.27 \times 10^{-1}$ | **+/100** |
| Effect: GR | $1.43 \times 10^{-2}$ | $-4.87 \times 10^{-2}$ | $7.62 \times 10^{-2}$ | |
| Effect: LAB | $-1.21 \times 10^{-1}$ | $-1.86 \times 10^{-1}$ | $-5.91 \times 10^{-2}$ | **−/100** |
| Effect: M1 | $-7.66 \times 10^{-2}$ | $-1.40 \times 10^{-1}$ | $-1.36 \times 10^{-2}$ | **−/99.0** |
| Effect: M2 | $-9.63 \times 10^{-2}$ | $-1.66 \times 10^{-1}$ | $-2.61 \times 10^{-2}$ | **−/99.6** |
| Effect: UT | $-7.13 \times 10^{-2}$ | $-1.30 \times 10^{-1}$ | $-9.49 \times 10^{-3}$ | **−/98.8** |
| Effect: SW | $1.16 \times 10^{-1}$ | $3.85 \times 10^{-2}$ | $1.93 \times 10^{-1}$ | **+/99.8** |
| Recipient: egg number |  |  |  |  |
|  | Mean | Lower 2.5% CI | Upper 2.5% CI | Effect/PD |
| $\beta 1$: intercept | 5.43 | 5.26 | 5.60 | |
| $\beta 2$: size | $2.47 \times 10^{-1}$ | $1.05 \times 10^{-1}$ | $3.96 \times 10^{-1}$ | **+/100** |
| $\beta 3$: age | $-1.47 \times 10^{-1}$ | $-2.82 \times 10^{-1}$ | $-1.03 \times 10^{-2}$ | **−/98.2** |
| $\beta 4$: mate | $-4.38 \times 10^{-2}$ | $-2.10 \times 10^{-1}$ | $1.20 \times 10^{-1}$ | |
| $\beta 5$: egg size | $-1.17 \times 10^{-1}$ | $-2.26 \times 10^{-1}$ | $-7.14 \times 10^{-3}$ | **−/98.5** |
| Effect: AU | $1.10 \times 10^{-1}$ | $-8.12 \times 10^{-2}$ | $3.19 \times 10^{-1}$ | |
| Effect: BE | $-1.93 \times 10^{-1}$ | $-4.63 \times 10^{-1}$ | $4.73 \times 10^{-2}$ | |
| Effect: FI | $1.98 \times 10^{-1}$ | $-8.48 \times 10^{-2}$ | $5.73 \times 10^{-1}$ | |
| Effect: FL | $1.48 \times 10^{-2}$ | $-1.83 \times 10^{-1}$ | $2.16 \times 10^{-1}$ | |
| Effect: FR | $1.99 \times 10^{-1}$ | $-1.50 \times 10^{-2}$ | $4.40 \times 10^{-1}$ | |
| Effect: GR | $1.47 \times 10^{-1}$ | $-4.77 \times 10^{-2}$ | $3.68 \times 10^{-1}$ | |
| Effect: LAB | $-2.17 \times 10^{-3}$ | $-2.26 \times 10^{-1}$ | $2.12 \times 10^{-1}$ | |
| Effect: M1 | $-1.81 \times 10^{-1}$ | $-4.37 \times 10^{-1}$ | $4.78 \times 10^{-2}$ | |
| Effect: M2 | $-2.23 \times 10^{-2}$ | $-2.73 \times 10^{-1}$ | $2.03 \times 10^{-1}$ | |
| Effect: UT | $8.86 \times 10^{-2}$ | $-1.10 \times 10^{-1}$ | $2.91 \times 10^{-1}$ | |
| Effect: SW | $-3.59 \times 10^{-1}$ | $-6.60 \times 10^{-1}$ | $-4.82 \times 10^{-2}$ | **−/98.9** |

### Phylogenetic relationship and signals

The estimated distance-based phylogeny represented the sampled geographic locations well (Fig. S1, Table S3). Specifically, western European populations (BE/FR/FL/UT/LAB/M1/M2) formed a monophyletic group, while a distinct monophyletic group was observed for other European populations (FI/GR/SW). French (FR/FL) and Dutch (UT/LAB/M1/M2) populations formed different monophyletic groups, and furthermore the neighbouring M1 and M2 populations were the most closely related. However, we did not detect any significant phylogenetic signals using mating success rates, PC1s, egg size and egg number (Table 6).

**Table 6.** Results of phylogenetic signals for each reproductive trait.

|  | Donor: male PC1 |  | Donor: female PC1 |  | Recipient: male PC1 |  | Recipient: female PC1 |  |
| --- | --- | --- | --- | --- | --- | --- | --- | --- |
|  | value | *p* | value | *p* | value | *p* | value | *p* |
| $C_{\text{mean}}$ | $-1.50 \times 10^{-1}$ | 0.542 | $-3.57 \times 10^{-2}$ | 0.305 | $-1.29 \times 10^{-1}$ | 0.501 | $8.01 \times 10^{-2}$ | 0.142 |
| $I$ | $-9.95 \times 10^{-2}$ | 0.469 | $-8.37 \times 10^{-2}$ | 0.131 | $-9.23 \times 10^{-2}$ | 0.260 | $-9.23 \times 10^{-2}$ | 0.224 |
| $K$ | $9.01 \times 10^{-1}$ | 0.601 | 1.04 | 0.131 | 1.02 | 0.280 | $9.74 \times 10^{-1}$ | 0.33 |
|  | Donor: mating success rate |  | Recipient: mating success rate |  | Average egg size |  | Total egg number |  |
|  | value | *p* | value | *p* | value | *p* | value | *p* |
| $C_{\text{mean}}$ | $-2.78 \times 10^{-1}$ | 0.849 | $-2.96 \times 10^{-1}$ | 0.833 | $2.74 \times 10^{-1}$ | 0.051 | $-1.69 \times 10^{-1}$ | 0.643 |
| $I$ | $-1.19 \times 10^{-1}$ | 0.929 | $-1.11 \times 10^{-1}$ | 0.807 | $-8.72 \times 10^{-2}$ | 0.157 | $-9.85 \times 10^{-2}$ | 0.470 |
| $K$ | $7.54 \times 10^{-1}$ | 0.945 | $8.52 \times 10^{-1}$ | 0.755 | 1.01 | 0.218 | $9.62 \times 10^{-1}$ | 0.388 |

### Discussions

### Diversity of reproductive traits among populations

Our observations and modelling clearly reveal diversity in reproductive traits among *Lymnaea stagnalis* populations, encompassing behaviours and reproductive investment in egg laying. Since these populations have been maintained for multiple generations under standardised laboratory conditions, this divergence in reproductive traits likely results from the evolutionary background of each population. In particular, the BE and M2 populations exhibited opposite effects on many examined characteristics. Namely, the BE population seems to be active in male behaviours (notably circling [C]; positioning [PS]; probing [PR]; eversion [E]; and failed insemination [INx]) as a recipient and in female behaviours (notably crawl out [CO/CO+]; and floating[F+]) as a donor, and is more successful at inseminating as an assigned recipient (i.e. mating success after 4d-isolation). In other words, this population seems to be more motivated to mate in the male role then what has previously been reported for our lab population (respectively, this study and Koene and Ter Maat 2007) and as a consequence also expresses more female behaviours that have been suggested to be indications of insemination avoidance (e.g. Palmeira et al. 2023, Daupagne and Koene 2020). In contrast, the M2 population seems to be rather inactive as a recipient and in female behaviours as a donor, and exhibits lower mating success in both roles, which may reflect lower motivation to mate in general. This tendency toward either active or inactive behaviours extends to some other populations (e.g. active: LAB and GR; inactive: UT and SW). The behaviours that consistently cluster together in the principal component analysis clearly indicate that the sperm donor behaviours are part of a stereotypical sequence of behaviours leading up to insemination that are known to generally occur in the order of circling, positioning, probing, eversion and intromission (as has been reported many times before). Interestingly, mounting does not cluster together with these behavioural components, indicating that this also occurs at moments that are unrelated to attempts to inseminate. The same applies to sperm recipient behaviours like crawl-out and floating, although the data show this less prominently.

The consistent tendencies in reproductive behaviours between population that we report here might be associated with mating motivation. In *L. stagnalis* previous studies that reported synchronous changes in mating behaviours in response to experimental manipulations suggest that these should be regarded as changes in male or female mating motivation (Moussaoui et al. 2018; Daupagne and Koene 2020). Moreover, mating motivation may vary between populations, which could be a consequence of sexual selection and sexual conflict (Moussaoui et al. 2018; Daupagne and Koene 2020; Palmeira et al. 2023). Further clarification of this issue is difficult in our study due to the experimental design, because to demonstrate that consistent motivational differences are present between and within populations would require repeated quantification of mating behaviour of the same individuals with different individuals.

The total egg numbers during 7 days laid varied significantly across populations, with the least productive population producing nearly 150 eggs less than the most productive one. In contrast, egg size remained relatively consistent across populations with the largest population difference being around 0.3 mm^2^. In addition, our modeling indicated that a higher egg number negatively correlated with egg size and vice versa, suggesting a certain trade-off between these traits, although a trade-off model incorporating total resource availability and its allocation could not be employed. Variation of egg (or seed) number and the trade-off with size are very well documented pattern across many organisms (Fleming and Gross 1990; Berrigan 1991; Christians 2000; Charnov and Ernest 2006; Sadras 2007). The number and size of eggs, often interpreted as proxies for reproductive quantity and quality, as well as the degree of resource allocation, have profound implications for fitness and are therefore subject to evolutionary processes driven by natural selection (Smith and Fretwell 1974). Notably, high variability in egg number and relative stability in egg size have been reported in taxa where parents provide little or no parental care to their offspring such as *L. stagnalis* (Smith and Fretwell 1974; Ebert 1994; Sadras 2007). In contrast, our modeling estimated that the effect of each population was more pronounced for egg size, with few significant effects on egg number. At first glance, this pattern appears to contradict the previous suggestion— high variability in egg number and stability in egg size. However, it is presumed that the variation in egg number can be strongly influenced by phenotypic plasticity in general (Sadras 2007), which may obscure the population effects. This hypothesis is further supported by the significant effects of both body size and age on egg number in our model. Significant variation in egg number was reported among geographically adjacent *L. stagnalis* populations (Puurtinen et al. 2007), and variability in egg number among populations might represent a fundamental character shared among *L. stagnalis*.

### Evolutionary insights

No phylogenetic signals were detected for the diversity of reproductive traits observed in this study. This suggests that the diversified traits examined here did not evolve along a branch of the phylogenetic tree under the Brownian motion model. However, the limited number of populations to compare may reduce the statistical power to detect such phylogenetic signals (Münkemüller et al. 2012). Notable differences in mating success were observed between the genetically (and geographically) closely related M1 and M2 populations, and the LAB population exhibited reproductive behaviours that contrasted markedly with those of other genetically-related Dutch populations (i.e. M1, M2, UT). These patterns support the statistical lack of a phylogenetic signal and rather suggest that reproductive traits of *L. stagnalis* may not be strongly constrained by phylogenetic lineage.

It is important to note, however, that while the Brownian motion-based evolutionary model was rejected, other evolutionary processes remain possible. For instance, rapid evolution driven by divergent selection pressures or founder effect could be potential evolutionary hypotheses. The significant differences between the closely related M1 and M2 populations, the distinctiveness of the LAB population, and the previously observed diversity in egg number among populations within the same drainage system (Puurtinen et al. 2007) can all be explained by rapid adaptation and/or founder effects. In freshwater snails, founder effects and rapid evolution are frequently reported (e.g. Angers et al. 2003; Pfenninger et al. 2011; Chapuis et al. 2017; Kagawa et al. 2019), and in *L. stagnalis* populations appear to be strongly influenced by genetic bottlenecks and drift (Kopp et al. 2012). Future research involving comprehensive multi-population analyses may provide further validation of these evolutionary patterns and offer deeper insights into the mechanisms driving the diversification of reproductive traits in this species and a full analysis of the lcWGS data may also be revealing here.

### Comparison to other studies

The differences observed in this study reflect responses in a controlled laboratory environment, and it remains uncertain whether the same trends would be observed in different environmental contexts. Indeed, environmental differences in laboratory conditions can have a variety of effects on behaviours (Griffith et al. 2017). This differential response to environments is likely attributable to phenotypic plasticity, but this plasticity is somehow shaped by evolutionary background (Miner et al. 2005; Pfening 2021). Interestingly, no significant population effects on male behaviour PCs were detected under the 7 days isolation treatment (i.e., designed to enhance male motivation), nor were there significant population effects estimated on female behaviours under the 4 days isolation treatment. These findings suggest that the prior treatments may have standardised the behavioural differences among populations. This indicates that carefully designed treatments before the mating experiments can be valuable to compare the differences focused on.

Several traits examined in this study have been the focus of previous research on *L. stagnalis*. However, differences in experimental settings, conditions, and *L. stagnalis* populations used in these studies usually make direct comparisons challenging. Nevertheless, certain parallels can be drawn. For example, our findings revealed a positive correlation between size and egg number, and the same relationship was reported in a previous study (Koene et al. 2007). Also, our results align with prior observations of a negative relationship between increasing age and male mating success (Hermann et al. 2009). It should be noted that the present analysis did not incorporate age- or size-population interactions in order to avoid overcomplicating the model. Nonetheless, the observed increase in egg number with larger body size and the decline in male mating success with ageing seems to represent general tendencies of *L. stagnalis*, rather than a specific feature of individual populations.

Interestingly, the presence or absence of mating had no statistically significant effect on either egg size and number in our models, although reduction of egg production after mating was documented in previous studies on *L. stagnalis* (Koene et al. 2009, 2010; Nakadera et al. 2014b; Swart et al. 2020). The primary effect of mating revealed by these studies is a short-term (less than a week) suppression in egg production, mainly driven by accessory gland proteins (ACPs: Koene et al. 2010; Nakadera et al. 2014b). Over the long term (more than a week), however, a low mating rate is likely to result in an increase in egg production compared to self-fertilisation (Koene et al. 2006; Hoffer et al. 2012, 2017). The discordance between our findings and previous studies could be explained by several factors. First, since only egg data after a week was collected in our study, the short-term suppression by ACPs may have already diminished, while any positive effect of mating could have become apparent at the 7^th^ day. Supporting this notion, a prior multi-population study found no significant effect of mating on egg production over a two-week period (Puurtinen et al. 2007).

Second, it is possible that each population represents a different tendency regarding the mating effect. Due to limitations in sample size and the need to avoid excessive model complexity, we did not construct models incorporating an interaction between population and mating status. Consequently, we were unable to assess mating effects that may occur in opposing directions across populations. In fact, when comparing egg size, number and production across the populations with and without mating, the LAB population—used in several prior studies—showed reduced egg outputs in mated individuals, whereas some populations showed increased outputs under the same conditions (Fig. S2). To elucidate the population-level differences in the mating effects, further experiments focused on each population are needed and it could provide important insights into the reproductive biology in *L. stagnalis*.

Our findings and modelling clearly highlighted substantial variations in reproductive traits among populations. This insight underscores the importance of avoiding assumptions of uniformity in species reproductive traits. Furthermore, many behavioural and physiological experiments rely on a single laboratory strain, which limits the generalisability of their findings to the entire species. While conducting multi-population experiments is often logistically challenging, uncovering the diversity and variability of reproductive traits within species—particularly those associated with reproductive isolation—could provide critical insights into the evolutionary and ecological dynamics underlying species differentiation.

## Acknowledgements

We are grateful to Marie-Agnés Coutellec, Otto Seppälla, Anssi Karvonen, Alexandra Staikou, Marianthi Hatziioannou, Jukka Jokela, and Tom van den Neucker for helping with collecting these European populations of *Lymnaea stagnalis*. We also thank Omar Bellaoui for maintaining the strains at Vrije University Amsterdam, and Yasuto Ishii for helpful comments on genomic distance analysis. The genetic data using lcWGS is part of comprehensive project for population genetics by JMK and YN.

## Author Contributions

Susanne Pekelharing (Conceptualization [equal], Investigation [lead], Methodology [support], Data curation [lead], Formal analysis [support], Writing—original draft [equal], Writing—review & editing [equal]), Yumi Nakadera (Data curation [support], Investigation [support], Methodology [equal], Resource [equal], Writing—review & editing [equal], Supervision [support], Funding acquisition [support]), Takumi Saito (Conceptualization [support], Investigation [support], Data curation [support], Formal analysis [lead], Visualization [lead], Writing—original draft [lead], Writing—review & editing [equal], Supervision [support], Funding acquisition [support]), and Joris M. Koene (Conceptualization [equal], Methodology [equal], Resource [equal], Writing—review & editing [equal], Supervision [lead], Project administration [lead], Funding acquisition [lead])

## Data accessibility statement

All data generated during this study are included in this published article, its supplementary materials, Yoda Portal by VU Amsterdam and deposited in GenBank.

---

**Figure S1.**
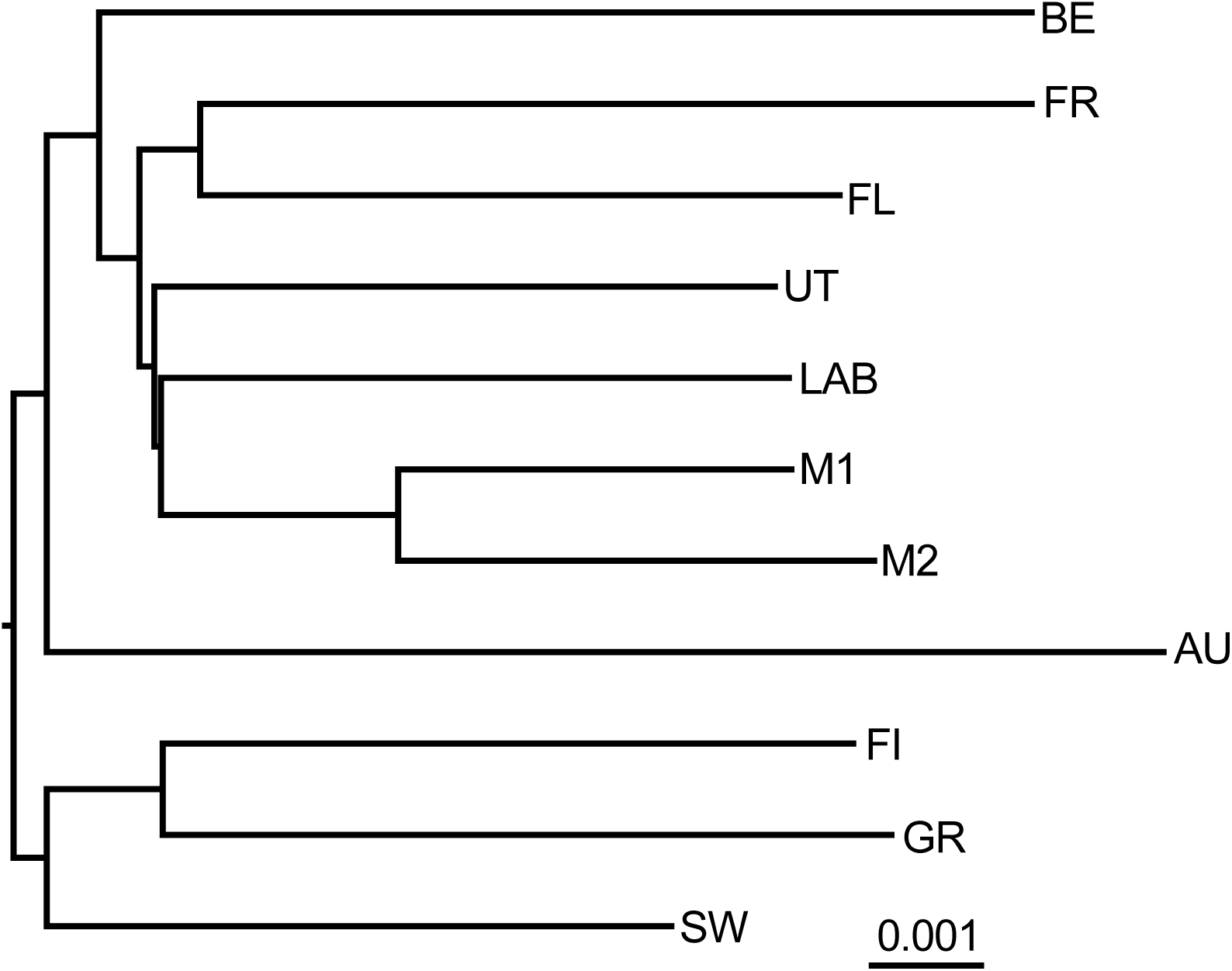
Distance based taxon addition tree with minimum-evolution principle using low-coverage whole genome sequencing. Codes indicate populations and raw distances are shown in Table S2.

**Figure S2.**
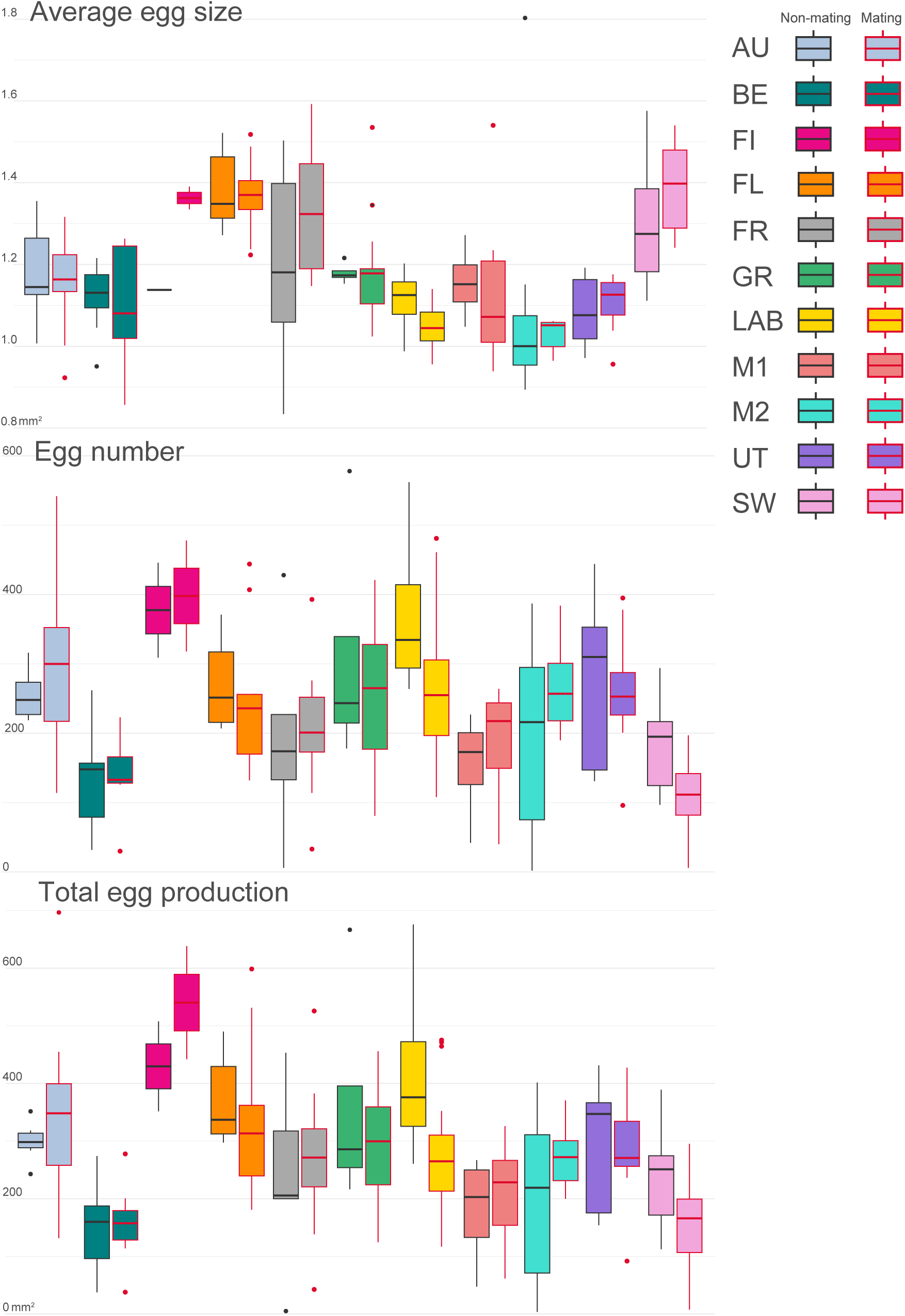
Box plots for egg size, number and production of each population. Codes and colours indicate populations (Table 1) and red enclosures indicate the presence of mating.

